# Network Analysis Prioritizes DEWAX and ICE1 as the Candidate Genes for Major eQTL Hotspots in Seed Germination

**DOI:** 10.1101/2020.04.29.050567

**Authors:** Margi Hartanto, Ronny V. L. Joosen, Basten L. Snoek, Leo A. J. Willems, Mark G. Sterken, Dick de Ridder, Henk W. M. Hilhorst, Wilco Ligterink, Harm Nijveen

## Abstract

Seed germination is characterized by a constant change of gene expression across different time points. These changes are related to specific processes, which eventually determine the onset of seed germination. To get a better understanding on the regulation of gene expression during seed germination, we performed a quantitative trait locus mapping of gene expression (eQTL) at four important seed germination stages (primary dormant, after-ripened, six-hour after imbibition, and radicle protrusion stage) using *Arabidopsis thaliana* Bay x Sha recombinant inbred lines (RILs). The mapping displayed the distinctness of the eQTL landscape for each stage. We found several eQTL hotspots across stages associated with the regulation of expression of a large number of genes. Interestingly, an eQTL hotspot on chromosome five collocates with hotspots for phenotypic and metabolic QTLs in the same population. Finally, we constructed a gene co-expression network to prioritize the regulatory genes for two major eQTL hotspots. The network analysis prioritizes transcription factors DEWAX and ICE1 as the most likely regulatory genes for the hotspot. Together, we have revealed that the genetic regulation of gene expression is dynamic along the course of seed germination.

**One-sentence summary:** Two transcription factors, DEWAX and ICE1, may be important regulators of gene expression during seed germination, based on network analysis of eQTL hotspots.

## Introduction

Seed germination involves a series of events starting with the transition of *quiescent* to physiologically active seeds and ends with the emergence of the embryo from its surrounding tissues. Germination is initiated when seeds become imbibed by water, leading to the activation of seed physiological activities (Nonogaki et al., 2010; Bewley et al., 2013). Major metabolic activities occur after seeds become hydrated, for example, restoration of structural integrity, mitochondrial repair, initiation of respiration, and DNA repair (Nonogaki et al., 2010; Bewley et al., 2013). For some species such as *Arabidopsis thaliana*, germination can be blocked by seed dormancy. Dormant seeds need to sense and respond to environmental cues to break their dormancy and complete germination. In *Arabidopsis thaliana*, seed dormancy can be alleviated by periods of dry after-ripening or moist chilling (Bewley et al., 2013). Soon after dormancy is broken, the storage reserves are broken down, and germination-associated proteins are synthesized. Lastly, further water uptake followed by cell expansion leads to radicle protrusion through endosperm and seed coat, which marks the end of germination (Bewley et al., 2013).

A major determinant for the completion of seed germination is the transcription and translation of mRNAs. The activity of mRNA transcription is low in dry, mature seeds (Comai and Harada, 1990; Leubner-Metzger, 2005), and drastically increases after seeds become rehydrated (Bewley et al., 2013). Nevertheless, stored mRNAs of more than 12,000 genes with various functions are already present in dry seeds. These mRNAs are not only remnants from the seed developmental process, but also mRNAs for genes related to metabolism as well as protein synthesis and degradation required in early seed germination (Rajjou et al., 2004; Nakabayashi et al., 2005). Later in after-ripened seeds, only a slight change in transcript composition was detected compared to the dry seeds (Finch-Savage et al., 2007). The major shift in transcriptome takes place after water imbibition (Nakabayashi et al., 2005). Interestingly, the transcriptome at the imbibition stage depends on the status of dormancy. For non-dormant seeds, most of the transcripts are associated with protein synthesis, while for dormant seeds, the transcripts are dominated by genes associated with stress-responses (Finch-Savage et al., 2007; Buijs et al., 2019). Even the transcript composition in primary dormant seeds, which occurs when the dormancy is initiated during development, is different from that of secondary dormant seeds, which occurs when the dormancy is reinduced (Cadman et al., 2006). These findings show the occurrence of phase transitions in transcript composition along the course from dormant to germinated seed.

As omics technology becomes more widely available, several transcriptomics studies in seed germination processes have been conducted on a larger-scale. More developmental stages, i.e., stratification and seedling stage, and even spatial analyses have been included in these studies, resulting in the identification of gene co-expression patterns as well as the predicted functions of hub-genes (Bassel et al., 2011; Narsai et al., 2011; Dekkers et al., 2013; Silva et al., 2016). Through guilt-by-association, these co-expression based studies can be used for the identification of regulatory genes that are involved in controlling the expression of downstream genes. These regulatory genes can be subjected to further studies by reverse genetics to provide more insight into the molecular mechanisms of gene expression in seed germination (i.e., Silva et al., 2016). Nevertheless, this approach still has limitations. Uygun et al. (2016) argued that co-expressed genes do not always have similar biological functions. On the other hand, genes involved in the same function are not always co-expressed since gene expression regulation could be the result of post-transcriptional or other layers of regulation (Lelli et al., 2012). Further, Uygun et al. (2016) emphasized the importance of combining the expression data with multiple relevant datasets to maximize the effort in the prioritization of candidate regulatory genes.

Genetical genomics is a promising approach to study the regulation of gene expression by combining genome-wide expression data with genotypic data of a segregating population (Jansen and Nap, 2001). To enable this strategy, the location of markers associated with variation in gene expression is mapped on the genome, which results in the identification of expression quantitative trait loci (eQTLs). Relative to the location of the associated gene, the eQTL can be locally or distantly mapped, known as local and distant eQTLs (Brem et al., 2002; Rockman and Kruglyak, 2006). Local eQTLs mostly arise because of variations in the corresponding gene or a cis-regulatory element. In contrast, distant eQTLs typically occur due to polymorphism on trans-regulatory elements located far away from the target genes (Rockman and Kruglyak, 2006). Therefore, given the positional information of distant eQTLs, one can identify the possible regulators of gene expression. However, the eQTL interval typically spans a large area of the genome and harbors hundreds of candidate regulatory genes. A large number of candidate genes would cause the experimental validation (e.g. using knock-out or overexpression lines) to be costly and take a long time. Therefore, a prioritization method is needed to narrow down the list of candidate genes underlying eQTLs, particularly on distant eQTL hotspots. A distant eQTL hotspot is a genomic locus where a large number of distant eQTLs are collocated (Breitling et al., 2008). The common assumption is that the hotspot arises due to one or more polymorphic master regulatory genes affecting the expression of multiple target genes (Breitling et al., 2008). Therefore, the identification of master regulatory genes becomes the center of most genetical genomics studies as the findings might improve our understanding of the regulation of gene expression (i.e., in Keurentjes et al., 2007; Jimenez-Gomez et al., 2010; Terpstra et al., 2010; Valba et al., 2015; Sterken et al., 2017).

In this study, we carried out eQTL mapping to reveal loci controlling gene expression in seed germination. To capture whole transcriptome changes during seed germination, we included four important seed germination stages, which are primary dormant seeds (PD), after-ripened seeds (AR), six-hours imbibed seeds (IM), and seeds with radicle protrusion (RP). In total, 160 recombinant inbred lines (RILs) from a cross between genetically distant ecotypes Bay-0 and Shahdara (Bay x Sha) were used in this study (Loudet et al., 2002). Our results show that each seed germination stage has a unique eQTL landscape, confirming the stage-specificity of gene regulation, particularly for distant regulation. Based on network analysis, we identify the transcription factors ICE1 and DEWAX as prioritized candidate regulatory genes for two major eQTL hotspots in PD and RP, respectively. Finally, the resulting dataset complements the previous phenotypic QTL (Joosen et al., 2012) and metabolite QTL (Joosen et al., 2013) datasets, allowing systems genetics studies in seed germination. The identified eQTLs are available through the web-based AraQTL (http://www.bioinformatics.nl/AraQTL/) workbench (Nijveen et al., 2017).

## Results

### Major transcriptional shifts take place after water imbibition and radicle protrusion

To visualize the transcriptional states of the parental lines and the RILs at the four seed germination stages, we performed a principal component analysis using the log-intensities of all expressed genes (Figure 1). The first principal component explains 55.6% of the variation and separates the samples into three groups. Germination progresses from left to right with the PD and AR seeds grouping together, indicating that the after-ripening treatment does not induce a considerable change in global transcript abundance. The large-scale transcriptome change only happens after water imbibition and radicle protrusion. This event was also observed by Finch-Savage et al. (2007) and Silva et al. (2016). The second principal component on the PCA explains 14.2% variance in the data and separates the RILs within each of the three clusters but not the parents. The source of this variation may be the genetic variation among samples and shows transgressive segregation of gene expression in RILs due to genetic reshuffling of the parental genomes during crossing and generations of selfing.

**Figure 1.** Principal component plot derived from transcriptome measurements of 164 RILs, and the Bay-0 and Sha parental lines taken at primary dormant seed (PD), after-ripened seed (AR), six-hours after imbibition (IM), and at the time when the radicle is protruded (RP).

To identify specific expression patterns among genes in the course of seed germination, we performed an additional analysis of the transcriptome data using hierarchical clustering (Figure 2). For this analysis, we only selected the 990 genes with a minimal fold change of two between any two consecutive stages (PD to AR, AR to IM, IM to RP). We then clustered both the genes and the seed samples. As shown in the figure, the clustering of samples shows similar grouping as in the previous PCA plot; three clusters were formed with one cluster containing both PD and AR, while IM and RP form separate clusters.

**Figure 2.** Hierarchical clustering of Bay-0, Sha, and 164 RILs transcriptome samples measured at four different seed germination stages (top) and 990 genes differentially expressed between two consecutive stages (left). Listed genes are the sample of genes for each cluster. Some enriched gene ontology terms for gene clusters are listed on the right.

The clustering of genes shows at least three distinctive gene expression patterns. In the first pattern, transcript abundance is highest in the last stage, radicle protrusion. A GO enrichment test suggests that transcripts with this expression pattern are involved in the transition from the heterotrophic seed to the autotrophic seedling stage, with enriched processes such as photosynthesis, response to various light, and response to temperature. This is in agreement with Rajjou et al. (2004), who showed that genes required for seedling growth are expressed after water imbibition. The second pattern shows an opposite trend with higher transcript abundances in the first three stages and lower expression at the end of the seed germination process. Some of these transcripts may be the remnant of seed development since the GO term related to this process is overrepresented. Moreover, transcripts involved in response to hydrogen peroxide were also overrepresented, which provides more evidence for the importance of reactive oxygen species in seed germination (for review see Wojtyla et al., 2016). The last pattern represents genes that are upregulated at the IM stage. Genes with this pattern are functionally enriched in the catabolism of fatty acids, a likely source of energy for seedling growth (Bewley et al., 2013). Altogether, these results suggest that co-expression patterns of genes reflect particular functions during the seed germination process.

### Distant eQTLs explain less variance than local eQTLs and are more specific to a seed germination stage

To map loci associated with gene expression levels, we performed eQTL mapping of 29,913 genes for each seed population representing four seed germination stages (Table 1). We found eQTLs, numbers ranging from 1,335 to 1,719 per stage (FDR = 0.05), spread across the genome. Among the genes with an eQTL, only a few (less than 1%) had more than one. We then categorized the eQTLs into local and distant based on the distance between the target gene and the eQTL peak marker or the confidence interval. Based on this criterion, over 72% of the eQTLs per stage were categorized as local, while the remainder were distant. Although the total of the identified eQTLs was different between the stages, the ratio of distant to local eQTLs was relatively similar for all stages. We then calculated the fraction of the total variation that is explained by the simple linear regression model for each eQTL. By comparing the density distributions (Figure S1), we showed that local eQTLs generally explain a more substantial fraction of gene expression variation than distant eQTLs. Finally, we determined the number of specific and shared eQTLs across stages (Figure 3). Here, we show that distant eQTLs are more specific to seed germination stages. Local eQTLs, on the other hand, are commonly shared between two or more stages, which is in line with previous experiments showing overlapping local eQTLs and specific distant eQTLs across different developmental stages (Vinuela et al., 2010), environments (Snoek et al., 2012; Lowry et al., 2013; Snoek et al., 2017) and populations (Cubillos et al., 2012).

**Table 1.** Summary of the eQTL mapping for the four different seed germination stages

| stage | eQTLs | genes with an eQTL | eQTL type | total | proportion |
| --- | --- | --- | --- | --- | --- |
| **primary dormant** | 1,335 | 1,328 | local | 955 | 0.72 |
|  |  |  | distant | 380 | 0.28 |
| **after-ripened** | 1,395 | 1,377 | local | 1,089 | 0.78 |
|  |  |  | distant | 306 | 0.22 |
| **six hours after imbibition** | 1,719 | 1,702 | local | 1,320 | 0.77 |
|  |  |  | distant | 399 | 0.23 |
| **radicle protrusion** | 1,426 | 1,418 | local | 1,096 | 0.77 |
|  |  |  | distant | 330 | 0.23 |

**Figure 3.** Shared local and distant eQTLs per seed germination stage.

### An eQTL hotspot on chromosome 5 is associated with genes related to seed germination and collocates with multiple metabolic and phenotypic QTLs

To get an overview of how the eQTLs were mapped over the genome, we visualized the eQTL locations and their associated genes on a local/distant eQTL plot (Figure 4A). Here, the local eQTLs are aligned across the diagonal and spread relatively equally across the genome, while it is not the case for the distant eQTLs. Furthermore, specific loci show clustering of eQTLs, which could indicate the presence of major regulatory genes that cause genome-wide gene expression changes. We identified ten so-called (distant-) eQTL hotspots, with at least two hotspots per stage (Table 2). The number of distant eQTLs located within these hotspots ranges from 16 to 96. The major eQTL hotspots are PD2, IM2, and RP4, with 69, 69, and 96 distant eQTLs co-locating, respectively. Moreover, the landscape of the eQTL hotspots (Figure 4B) differs for every stage, including PD and AR, which is surprising since these two stages have a relatively similar transcriptome profile (Figure 1).

**Table 2.** Distant eQTL hotspots of the four seed germination stages. These hotspots were identified by dividing the genome into bins of 2 Mbp and performing a test to determine whether the number of distant eQTLs on a particular bin is higher than expected (p > 0.0001) assuming a Poisson distribution. Seed germination phenotype and metabolite data were taken from Joosen et al. (2012) and Joosen et al. (2013), respectively. Detailed information about enriched GO terms, metabolite, and phenotype can be seen on Supplemental Table S2 in the Supplementary Material.

| hotspot ID | position | distant eQTLs | enriched GO terms | metabolite QTL | phenotype QTL |
| --- | --- | --- | --- | --- | --- |
| **PD1** | ch1:6-10 Mb | 43 | 11 | 1 | 4 |
| **PD2** | ch3:8-12 Mb | 69 | 3 | 2 | 1 |
| **AR1** | ch2:12-14 Mb | 16 | 0 | 0 | 0 |
| **AR2** | ch3:2-4 Mb | 20 | 9 | 1 | 1 |
| **IM1** | ch5:6-8 Mb | 19 | 2 | 24 | 1 |
| **IM2** | ch5:22-26 Mb | 69 | 6 | 6 | 31 |
| **RP1** | ch1:0-2 Mb | 23 | 1 | 0 | 1 |
| **RP2** | ch1:6-8 Mb | 18 | 0 | 0 | 3 |
| **RP3** | ch5:14-16 Mb | 21 | 29 | 0 | 1 |
| **RP4** | ch5:24-26Mb | 96 | 18 | 20 | 25 |

**Figure 4.** eQTL mapping from four different seed germination stages. The local-distant eQTL plot is shown on top (**A**). The positions of eQTLs are plotted along the five chromosomes on the x-axis and the location of the genes with an eQTL is plotted on the y-axis. The black dots (●) represent local eQTLs (located within 1 Mb of the associated gene) and the colored dots represent distant eQTLs (located far from the associated gene). The gray horizontal lines next to each dot indicate the confidence interval of the eQTL location based on a 1.5 drop in - log10(p-value). The histogram of the number of eQTLs per genomic location is shown at the bottom (**B**). The horizontal dashed black lines mark the significance threshold for an eQTL hotspot.

We remapped the QTLs for previously studied seed germination phenotypes (Joosen et al., 2012) and metabolites (Joosen et al., 2013) using the RNA-seq based genetic map (Serin et al., 2017). We then visualized the resulting QTL count histograms alongside the eQTL histogram (Figure 5). The histogram shows that several eQTL hotspots collocate with hotspots for phenotype and metabolite QTLs (phQTLs and mQTLs, respectively). The most striking example is the collocation of QTLs on chromosome 5 around 24—25 Mb (IM2 and RP4) at the last two stages of seed germination. We performed gene ontology (GO) term enrichment analysis for genes with an eQTL mapping to these hotspots, and found ‘seed germination’ enriched among other terms (Table 2). These findings taken together indicate that the IM2 and RP4 hotspots harbor one or more important genes affecting gene expression during seed germination. Therefore, the identification of the regulatory gene(s) for one of these hotspots can give us more insight into the trans-regulation of gene expression during seed germination.

**Figure 5.** Hotspots for phQTLs, mQTLs, and eQTLs. A region of interest is located on chromosome 5 (around 24—26 Mb) where hotspots from different QTL levels collocate.

### Transcription factors were prioritized as the candidate genes for major eQTL hotspots

To prioritize the candidate regulatory genes underlying eQTL hotspots in this study, we constructed a network based on the expression of genes with eQTLs on the hotspot location. We built the network for two hotspots: RP4, where QTLs for expression, metabolite, and phenotype are collocated; and PD2, another major eQTL hotspot in this study. For RP4, the total number of genes used to construct the network was 116, of which 20 had a local eQTL at the hotspot, whereas for PD2, 114 genes were identified, of which 45 with a local eQTL. The genes with local eQTLs were then labeled as candidates. The networks were constructed by integrating predictions from several gene regulatory network inference methods to ensure the robustness of the result (Marbach et al., 2012). The direction of the edges in the network is predicted using the GENIE3 method (Huynh-Thu et al., 2010). For each candidate gene, we calculated the outdegree, indicating the number of outgoing edges of a gene to other genes in the network, and the closeness centrality of the candidate gene nodes, which shows the efficiency of the gene in spreading information to the rest of the genes in the network (Pavlopoulos et al., 2011). Finally, these two network properties were used to prioritize the most likely regulator of the distant eQTL hotspot.

In the resulting network, genes encoding the transcription factors DECREASE WAX BIOSYNTHESIS/DEWAX (AT5G61590), and INDUCER OF CBP EXPRESSION 1/ICE1 (AT3G26744) were prioritized as the most likely candidate genes for RP4 (Figure 6) and PD2 (Figure 7), respectively. As many as 15 genes were predicted to be associated with DEWAX and 32 genes with ICE1. Note that these numbers depend on the chosen threshold; nonetheless, the current candidates are robust to changes when the parameter was changed (Table S3 and Table S4). Furthermore, these two genes also had the highest closeness centrality among the other candidates, showing that these genes have a strong influence within the network. We assessed the Bay x Sha SNP data (Genomes Consortium. Electronic address and Genomes, 2016) and found several SNPs between the Bay and Sha parents in both the DEWAX and ICE1 genes, including two that affect the amino acid sequence of the corresponding proteins (Table S5 and Table S6). Also, querying for DEWAX and ICE1 on AraQTL showed a local eQTL for both genes in an experiment using the same RIL population on leaf tissue (West et al., 2007). This evidence supports the presence of DEWAX and ICE1 polymorphisms between the Bay and Sha allele that might be responsible for the steadily occurring local eQTLs at three stages (PD, IM, RP) for DEWAX and all four stages for ICE1.

**Figure 6.** The prioritization of candidate genes for RP4 eQTL hotspot. The network of genes associated with RP4 is visualized in **A**. The genes in the network are represented by nodes with various sizes according to the outdegree. The unlabeled grey nodes are the targets (genes with a distant eQTL) and the labelled green nodes are the candidates (genes with a local eQTL). Nodes with a red border are transcription factors. The yellow node is DEWAX (AT5G61590). The list of top ten candidate genes for the hotspot is shown in **B**. The expression of DEWAX in 160 RILs across the four seed germination stages is visualized in **C**. The RILs with the Sha allele of the gene are depicted in blue, the ones with the Bay-0 allele of DEWAX are depicted in red.

**Figure 7.** The prioritization of candidate genes for the PD2 eQTL hotspot. The network of genes associated with PD2 is visualized in **A**. The genes in the network are represented by nodes with various sizes according to the outdegree. The unlabeled grey nodes are the targets (genes with a distant eQTL) and labelled green nodes are the candidates (genes with a local eQTL). Nodes with a red border are transcription factors. The yellow node is ICE1 (AT3G26744). The list of top ten candidate genes for the hotspot is shown in **B**. The expression of ICE1 in 160 RILs across the four seed germination stages is visualized in **C**. The RILs with the Sha allele of the gene are depicted in blue, the ones with the Bay-0 allele of ICE1 are depicted in red.

## Discussion

### The function of DEWAX may be related to seed cuticular wax biosynthesis

In this study, we constructed a network of genes associated with the RP4 eQTL hotspot and showed that *DEWAX* was prioritized as the candidate gene for the hotspot. *DEWAX* encodes an AP2/ERF-type transcription factor that is well-known as a negative regulator of cuticular wax biosynthesis (Go et al., 2014; Suh and Go, 2014; Cui et al., 2016; Li et al., 2019) and a positive regulator of defense response against biotic stress (Ju et al., 2017; Froschel et al., 2019). This gene also seems to be involved in drought stress response (Huang et al., 2008) by inducing the expression of genes that confer drought tolerance (Sun et al., 2016), some of which (*LEA4-5*, *LTI-78*) have a distant eQTL at the RP4 hotspot. Moreover, the overexpression of *DEWAX* in Arabidopsis increases the seed germination rate (Sun et al., 2016). The role of *DEWAX* in seed germination is still unknown but may be related to cuticular wax biosynthesis.

Wax is a mixture of hydrophobic lipids, which is part of the plant cuticle together with cutin and suberin (Yeats and Rose, 2013). Previous studies have demonstrated that the biosynthesis of wax in the cuticular layer of stems and leaves is negatively regulated by *DEWAX* (Go et al., 2014; Suh and Go, 2014; Cui et al., 2016; Li et al., 2019). Although the function of this gene has never been reported in seeds, the presence of a cuticular layer indeed plays a significant role in maintaining seed dormancy (De Giorgi et al., 2015; Nonogaki, 2019). In Arabidopsis seeds, the thick cuticular structure covering the endosperm prevents cell expansion and testa rupture that precede radicle protrusion. Besides, this layer also reduces the diffusion of oxygen into the seed, thus preventing oxidative stress that may cause rapid seed aging and loss of dormancy (De Giorgi et al., 2015).

Besides *DEWAX*, *MUM2* is another possible regulatory gene for the RP4 hotspot based on QTL confirmation of an imbibed seed size phenotype using a heterogeneous inbred family approach (Joosen et al., 2012). In our study, we also discovered that most eQTLs on the RP4 hotspot peak at the marker located closely to the MUM2 location (Figure S2), which provides more evidence for this gene as the regulator for the hotspot. *MUM2* encodes a cell-wall modifying beta-galactosidase involved in seed coat mucilage biosynthesis, and the *mum2* mutant is characterized by a failure in extruding mucilage after water imbibition (Dean et al., 2007). In our analysis, *MUM2* did not have a distant eQTL on the RP4 hotspot; thus, it is not prioritized as a prominent candidate, pointing out a limitation of our approach in prioritizing candidate eQTL hotspot genes which will be discussed later. Nonetheless, we found some evidence connecting *DEWAX* to *MUM2*. First, Shi et al. (2019) found out that the mutant of *CPL2*, another gene involved in wax biosynthesis, showed a delayed secretion of the enzyme encoded by *MUM2* that disrupts seed coat mucilage extrusion. In the same study, they revealed that *CPL2* encodes a phosphatase involved in secretory protein trafficking required for the secretion of extracellular matrix materials, including wax and cell wall-modifying enzyme. This finding provides a link between wax biosynthesis and cell-wall modifying enzymes, and possibly between the genes involved in these processes.

Second, the expression of *DEWAX* may be the consequence of the disruption of seed mucilage extrusion. Penfield et al. (2001) suggest that seed mucilage helps enhance water uptake to ensure efficient germination in the condition of low water potential. This is supported by the evidence that the mucilage-impaired mutant showed reduced maximum germination only on osmotic polyethylene glycol solutions (Penfield et al., 2001). Therefore, the absence of mucilage in imbibed seed under low water potential may cause osmotic stress in the seed and, in turn, induce the expression of DEWAX, which is known to play a role in the response of plants against osmotic stress (Sun et al., 2016). If this is the case, then a scenario could be that DEWAX acts downstream of MUM2, and the expression variation of these two genes lead to the emergence of the RP4 eQTL hotspot.

### Network analysis shows the involvement of ICE1 as a regulator of gene expression during seed germination

ICE1 is an MYC-like basic helix-loop-helix (bHLH) transcription factor that shows pleiotropic effects in plants. Earlier studies of ICE1 mostly focus on the protein function in the acquisition of cold tolerance (Chinnusamy et al., 2003; Lee et al., 2005) and stomatal lineage development (Kanaoka et al., 2008). Recently, ICE1 was also shown to form a heterodimer with ZOU, another bHLH transcription factor, to regulate endosperm breakdown required for embryo growth during seed development (Denay et al., 2014). At a later stage, ICE1 negatively regulates ABA-dependent pathways to promote seed germination and seedling establishment (Liang and Yang, 2015). This process involves repressing the expression of transcription factors in ABA signaling, such as ABI3 and ABI5, and ABA-responsive genes, such as *EM6* and *EM1*, thus initiating seed germination and subsequent seedling establishment (Hu et al., 2019; MacGregor et al., 2019).

In this study, we performed a network analysis for genes having distant eQTLs on the PD2 hotspot and prioritized ICE1 as the most likely regulator using network analysis. The high connectivity of ICE1 with the other genes in the network could reflect an essential regulatory function of this gene during seed germination. However, we did not find any of the known ICE1 target genes (i.e., *ABI3*, *ABI5*, *EM1*, and *EM6*) nor seed germination phenotype (Figure 5) having an eQTL at the *ICE1* locus. It could be that the ICE1 polymorphism is not severe enough to cause considerable trait variation, especially to break a robust biological system where several buffering mechanisms exist to prevent small molecular perturbation from propagating to the phenotypic level (Fu et al., 2009; Signor and Nuzhdin, 2018).

### Limitations of co-expression network in identifying candidate genes of eQTL hotspots

The construction of a co-expression network is a promising approach to prioritize candidate eQTL genes (Serin et al., 2016). Despite its potential, there is a major limitation in using a co-expression network. The network is based on gene expression data; hence the identified causal genes are those that directly affect gene expression. For example, as we described above, our approach did not prioritize *MUM2* for the RP4 hotspot, possibly because the gene does not cause variation in the target gene expression but rather causes differences at another level of target gene regulation (e.g., enzyme biosynthesis) between two parental alleles in the RIL population. Other studies reported similar results where a known causal gene was not detected as a hub in the network (Jimenez-Gomez et al., 2010; Sterken et al., 2017). To overcome this, future work should focus on networks that are built upon multi-omics data by including metabolic, proteomic, and, more importantly, phenotypic measurement data (Hawe et al., 2019). Moreover, prior biological knowledge, including protein-protein interaction (Szklarczyk et al., 2017), transcription factor binding-site (Kulkarni et al., 2018), and other types of interactions (for review see Kulkarni and Vandepoele, 2019) can be incorporated to construct data-driven interaction networks. Nevertheless, our approach offers a simple and straightforward way to prioritize candidate genes underlying eQTL hotspots from a limited amount of resources.

## Materials and Methods

### Plant materials

In this study, we used 164 recombinant inbred lines (RILs) derived from a cross between the Bay-0 and Shahdara Arabidopsis ecotypes (Loudet et al., 2002) provided by the Versailles Biological Resource Centre for Arabidopsis (http://dbsgap.versailles.inra.fr/vnat). The plants were sown in a fully randomized setup on 4×4 cm rockwool plugs (MM40/40, Groudan B. V.) and hydrated with 1 g/l Hyponex (NPK = 7:6:19, http://www.hyponex.co.jp) in a climate chamber (20°C day, 18°C night) with 16 hours of light (35 W/m2) at 70% relative humidity. Seeds from four to seven plants per RIL were bulk harvested for the experiment (see also Joosen et al., 2012; Joosen et al., 2013). The genotypic data consisting of 1,059 markers per line was obtained from Serin et al. (2017). However, the genotypic data is available only for 160 RILs; therefore, we used this number of lines for eQTL mapping.

### Experimental setup

The RIL population was grouped into four subpopulations, each one representing one of the four different seed germination stages. We used the designGG-package (Li et al., 2009) in R (version 3.6.0 Windows x64) to aid the grouping so that the distribution of Bay-0 and Sha alleles between sub-populations is optimized. The first stage is the primary dormant (PD) stage when the seeds were harvested and stored at −80°C after one week at ambient conditions. The second stage is after-ripened (AR) seeds that obtained maximum germination potential after five days of imbibition by storing at room temperature and ambient relative humidity. The third stage is the 6 hours imbibition (IM) stage. For this stage, the seeds were after-ripened and imbibed for six hours on water-saturated filter paper at 20°C and immediately transferred to a dry filter paper for 1 minute to remove the excess of water. The fourth stage is the radicle protrusion (RP) stage. To select seeds at this stage, we used a binocular to observe the presence of a protruded radicle tip.

### RNA isolation

Total RNA was extracted according to the hot borate protocol modified from Wan and Wilkins (1994). For each treatment, 20 mg of seeds were homogenized and mixed with 800 μl of extraction buffer (0.2M Na boratedecahydrate (Borax), 30 mM EGTA, 1% SDS, 1% Na deoxycholate (Na-DOC)) containing 1.6 mg DTT and 48 mg PVP40 which had been heated to 80°C. Then, 1 mg proteinase K was added to this suspension and incubated for 15 min at 42°C. After adding 64 μl of 2 M KCL, the samples were incubated on ice for 30 min and subsequently centrifuged for 20 min at 12,000 g. Ice-cold 8 M LiCl was added to the supernatant in a final concentration of 2 M, and the tubes were incubated overnight on ice. After centrifugation for 20 min at 12,000 g at 4°C, the pellets were washed with 750 μl ice-cold 2 M LiCl. The samples were centrifuged for another 10 min at 10,000 g at 4°C, and the pellets were re-suspended in 100 μl DEPC treated water. The samples were phenol-chloroform extracted, DNAse treated (RQ1 DNase, Promega), and further purified with RNeasy spin columns (Qiagen) following the manufacturer’s instructions. The RNA quality and concentration were assessed by agarose gel electrophoresis and UV spectrophotometry.

### Microarray analysis

RNA was processed for use on Affymetrix Arabidopsis SNPtile array (atSNPtilx520433), as described by the manufacturer. Briefly, 1 mg of total RNA was reverse transcribed using a T7-Oligo(dT) Promoter Primer in the first-strand cDNA synthesis reaction. Following RNase H-mediated second-strand cDNA synthesis, the double-stranded cDNA was purified and served as a template in the subsequent in vitro transcription reaction. The reaction was carried out in the presence of T7 RNA polymerase and a biotinylated nucleotide analog/ribonucleotide mix for complementary RNA (cRNA) amplification and biotin labeling. The biotinylated cRNA targets were then cleaned up, fragmented, and hybridized to the SNPtile array. The hybridization data were extracted using a custom R script with the help of an annotation-file based on TAIR10. Intensity data were log-transformed and normalized using the *normalizeBetweenArrays* function with the quantile method from Bioconductor package limma (Ritchie et al., 2015). Then, for each annotated gene, the log-intensities of anti-sense exon probes were averaged.

### Clustering analysis

Principal component analysis for log-intensities of all parents and RIL population samples was done using the pr.comp function in R where the unscaled log intensities are shifted to be zero centered. For hierarchical clustering, we only selected genes with a minimal fold change of 2 between any pair of consecutive stages (PD to AR, AR to IM, or IM to RP). Then, the distance matrices of filtered genes and all samples were calculated using the absolute Pearson correlation. These matrices were clustered using Ward’s method. We manually set the number of clusters to 8 and performed gene ontology enrichment for each of the clusters using the weight algorithm of the topGO package in R and used 29,913 genes detected by hybridization probes as the background (Alexa et al., 2006).

### eQTL mapping

For eQTL mapping, we used 160 RILs separated into four subpopulations, each representing one specific seed germination stage. For each stage separately, eQTLs were mapped using a single-marker model, as in Sterken et al. (2017). The gene expression data were fitted to the linear model

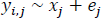

where y is the log-intensity representing the expression of a gene *i*(*i* = 1, 2, …, 29,913) of RIL *j*(*j* = 1, 2, …, 160) explained by the parental allele on marker location *x*(*x* = 1, 2, …, 1,059). The random error in the model is represented by *e_j_*.

To account for the multiple-testing burden in this analysis, we determined the genome-wide significant threshold using a permutation approach (e.g. see Sterken et al., 2017). A permuted dataset was created by randomly distributing the log-intensities of the gene under study over the genotypes. Then, the previous eQTL mapping model was performed on this permuted dataset. This procedure was repeated 100 times for each stage. The threshold was determined using:

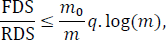

where, at a specific significance level, the false discoveries (FDS) were the averaged permutation result, and real discoveries (RDS) were the outcome of the eQTL mapping using the unpermuted dataset. The number of true hypotheses tested (*m*_0_) was 29,913 - RDS, and the number of hypotheses () tested was the number of genes, which was 29,913. For the -value, we used a threshold of 0.05. As a result, we got a threshold of 4.2 for PD and AR, 4.1 for IM, and 4.3 for RP.

The confidence interval of an eQTL was determined based on a -log_10_(*p*-value) drop of 1.5 compared to the peak marker (as in Keurentjes et al., 2007; Cubillos et al., 2012). We determine an eQTL as local if the peak marker or the confidence interval lies within 1 Mb or less from the target gene location (as in Cubillos et al., 2012). All eQTLs that did not meet this criterion were defined as distant.

We defined a region as an eQTL hotspot if the number of distant-eQTLs mapped to a particular genomic region significantly exceeded the expectation. First, we divided the genome into bins of 2 Mb. Then, we determined the expected number of distant-eQTLs per genomic bin by dividing the total number of distant-eQTLs by the total number of bins. Based on a Poisson distribution, any bin having an actual number of distant-eQTLs larger than expected (*p* < 0.0001) was then considered as an eQTL hotspot.

### Gene regulatory network inference and candidate genes prioritization of eQTL hotspot

We used a community-based approach to infer regulatory networks of genes with an eQTL on a hotspot location using expression data. In this approach, we assume the hotspot is caused by a polymorphism in or near one or more regulatory genes causing altered expression that can be detected as a local eQTL (Breitling et al., 2008; Joosen et al., 2009; Jimenez-Gomez et al., 2010; Serin et al., 2017). Based on this assumption, we labeled all genes with a local eQTL on a hotspot as candidate regulators and genes with a distant eQTL as targets. The expression of these genes was subjected to five different network inference methods to predict the interaction weight. The methods used were TIGRESS (Haury et al., 2012), Spearman correlation, CLR (Faith et al., 2007), ARACNE (Margolin et al., 2006), and GENIE3 (Huynh-Thu et al., 2010). The predictions from GENIE3 were used to establish the direction of the interaction by removing the one that has the lowest variable importance to the expression of the target genes between two pairs of genes. For instance, if the importance of gene_i_ – gene_j_ is smaller than gene_j_ – gene_i_, then the former is removed. By averaging the rank, the predictions of all inference methods were integrated to produce a robust and high performance prediction (Marbach et al., 2012). The threshold was determined as the minimum average rank where all nodes are included in the network. Finally, the network was visualized using Cytoscape (version 3.7.1) (Shannon et al., 2003), and network properties were calculated using the NetworkAnalyzer tool (Assenov et al., 2008). The candidate genes for each eQTL hotspot were prioritized based on their outdegree and closeness centrality (Pavlopoulos et al., 2011).

## Script availability

The code for the analysis and visualization is available in the form of R scripts at the Wageningen University GitLab repository (https://git.wur.nl/harta003/seed-germination-qtl).

## Accession numbers

Cel files of microarray data have been deposited in the ArrayExpress database at EMBL-EBI (www.ebi.ac.uk/arrayexpress) under accession number E-MTAB-xxxx.

## Supplemental materials

**Supplemental Figure S1.** Density distribution of the absolute eQTL effect, -log(p), and explained phenotypic variance (R2) for local and distant eQTLs.

**Supplemental Figure S2.** The histogram of the number of distant eQTLs per marker location for the PD2 (A) and RP4 (B) hotspot.

**Supplemental Table S1.** Gene ontology enrichment for genes with distinctive expression patterns during seed germination.

**Supplemental Table S2.** Distant eQTL hotspots of the four seed germination stages

**Supplemental Table S3.** The mean rank and standard deviation of candidate genes as the most likely causal genes for the RP4 hotspot across different thresholds

**Supplemental Table S4.** The mean rank and standard deviation of candidate genes as the most likely causal genes for the PD2 hotspot across different thresholds.

**Supplemental Table S5.** The location and type of SNPs on candidate genes for the RP4 eQTL hotspot and *MUM2.*

**Supplemental Table S6.** The location and type of SNPs on candidate genes for the PD2 eQTL hotspot.

**Supplemental Table S7.** The list of genetic markers used for QTL mapping.

**Supplemental Table S8.** The genetic map of Bay-0 x Sha parents and the RIL population.

**Supplemental Table S9.** Gene expression levels of Bay-0 x Sha parents and the RIL population.

**Supplemental Table S10.** Phenotype measurements of Bay-0 x Sha parents and the RIL population.

**Supplemental Table S11.** Metabolite measurements of Bay-0 x Sha parents and the RIL population.

**Supplemental Table S12.** Differentially expressed genes between any of two consecutive stages.

**Supplemental Table S13.** The list of expression QTL.

**Supplemental Table S14.** The list of phenotype QTL.

**Supplemental Table S15.** The list of metabolite QTL.

