## Supplemental Tables 1-6 and Supplemental Figures 1-2 for "Network Analysis Prioritizes DEWAX and ICE1 as the Candidate Genes for Major eQTL Hotspots in Seed Germination"

**Supplemental Table S1.** Gene ontology enrichment for genes with distinctive expression patterns during seed germination.

| Pattern | GO ID | Term | Annotated | Significant | Expected | Fisher | FDR | cluster |
|---|---|---|---|---|---|---|---|---|
| 1 | GO:0019684 | photosynthesis, light reaction | 271 | 45 | 4.02 | 1E-30 | 6E-27 | 3 |
| 1 | GO:0019682 | glyceraldehyde-3-phosphate metabolic pro... | 334 | 42 | 4.96 | 1E-26 | 2E-23 | 3 |
| 1 | GO:0042254 | ribosome biogenesis | 339 | 41 | 5.03 | 2E-25 | 3E-22 | 3 |
| 1 | GO:0015995 | chlorophyll biosynthetic process | 125 | 23 | 1.86 | 7E-19 | 8E-16 | 3 |
| 1 | GO:0006970 | response to osmotic stress | 833 | 48 | 12.37 | 6E-16 | 5E-13 | 3 |
| 1 | GO:0070838 | divalent metal ion transport | 209 | 24 | 3.1 | 9E-15 | 6E-12 | 3 |
| 1 | GO:0010256 | endomembrane system organization | 195 | 23 | 2.9 | 2E-14 | 1E-11 | 3 |
| 1 | GO:0006739 | NADP metabolic process | 186 | 23 | 2.76 | 5E-14 | 3E-11 | 3 |
| 1 | GO:0046686 | response to cadmium ion | 466 | 32 | 6.92 | 6E-13 | 3E-10 | 3 |
| 1 | GO:0009266 | response to temperature stimulus | 950 | 45 | 14.1 | 5E-12 | 2E-09 | 3 |
| 1 | GO:0009735 | response to cytokinin | 263 | 23 | 3.9 | 2E-11 | 6E-09 | 3 |
| 1 | GO:0010218 | response to far red light | 99 | 15 | 1.47 | 2E-11 | 7E-09 | 3 |
| 1 | GO:0019760 | glucosinolate metabolic process | 204 | 20 | 3.03 | 3E-11 | 1E-08 | 3 |
| 1 | GO:0010114 | response to red light | 104 | 15 | 1.54 | 4E-11 | 1E-08 | 3 |
| 1 | GO:0009637 | response to blue light | 128 | 15 | 1.9 | 8E-10 | 2E-07 | 3 |
| 1 | GO:0080147 | root hair cell development | 203 | 18 | 3.01 | 2E-09 | 4E-07 | 3 |
| 1 | GO:0009642 | response to light intensity | 282 | 21 | 4.19 | 2E-09 | 4E-07 | 3 |
| 1 | GO:0006833 | water transport | 141 | 15 | 2.09 | 3E-09 | 7E-07 | 3 |
| 1 | GO:0072330 | monocarboxylic acid biosynthetic process | 760 | 45 | 11.28 | 3E-09 | 7E-07 | 3 |
| 1 | GO:0009657 | plastid organization | 388 | 30 | 5.76 | 6E-09 | 1E-06 | 3 |
| 1 | GO:0055080 | cation homeostasis | 318 | 23 | 4.72 | 7E-09 | 1E-06 | 3 |
| 1 | GO:0080167 | response to karrikin | 128 | 14 | 1.9 | 7E-09 | 1E-06 | 3 |
| 1 | GO:0035304 | regulation of protein dephosphorylation | 140 | 14 | 2.08 | 2E-08 | 4E-06 | 3 |
| 1 | GO:0016126 | sterol biosynthetic process | 164 | 15 | 2.43 | 2E-08 | 4E-06 | 3 |
| 1 | GO:0016052 | carbohydrate catabolic process | 334 | 20 | 4.96 | 3E-08 | 5E-06 | 3 |
| 1 | GO:0042742 | defense response to bacterium | 398 | 23 | 5.91 | 3E-08 | 6E-06 | 3 |
| 1 | GO:0006084 | acetyl-CoA metabolic process | 85 | 11 | 1.26 | 5E-08 | 8E-06 | 3 |
| 1 | GO:0009744 | response to sucrose | 208 | 16 | 3.09 | 9E-08 | 1E-05 | 3 |
| 1 | GO:0016132 | brassinosteroid biosynthetic process | 118 | 12 | 1.75 | 2E-07 | 3E-05 | 3 |
| 1 | GO:0010155 | regulation of proton transport | 77 | 10 | 1.14 | 2E-07 | 3E-05 | 3 |
| 1 | GO:0009902 | chloroplast relocation | 104 | 11 | 1.54 | 4E-07 | 6E-05 | 3 |
| 1 | GO:0042335 | cuticle development | 49 | 8 | 0.73 | 6E-07 | 8E-05 | 3 |
| 1 | GO:0006094 | gluconeogenesis | 164 | 13 | 2.43 | 1E-06 | 1E-04 | 3 |
| 1 | GO:0006412 | translation | 495 | 23 | 7.35 | 2E-06 | 2E-04 | 3 |
| 1 | GO:0009825 | multidimensional cell growth | 111 | 11 | 1.65 | 2E-06 | 2E-04 | 3 |
| 1 | GO:0010196 | nonphotochemical quenching | 7 | 4 | 0.1 | 2E-06 | 2E-04 | 3 |
| 1 | GO:0071554 | cell wall organization or biogenesis | 880 | 33 | 13.07 | 2E-06 | 2E-04 | 3 |
| 1 | GO:0034284 | response to monosaccharide | 173 | 13 | 2.57 | 2E-06 | 2E-04 | 3 |
| 1 | GO:0006569 | tryptophan catabolic process | 79 | 9 | 1.17 | 3E-06 | 3E-04 | 3 |
| 1 | GO:0042737 | drug catabolic process | 137 | 11 | 2.03 | 7E-06 | 7E-04 | 3 |
| 1 | GO:0043481 | anthocyanin accumulation in tissues in r... | 113 | 10 | 1.68 | 7E-06 | 8E-04 | 3 |

| | | | | | | | | |
|---|---|---|---|---|---|---|---|---|
| 1 | GO:0019344 | cysteine biosynthetic process | 205 | 24 | 1.54 | 6E-22 | 3E-18 | 4 |
| 1 | GO:0010035 | response to inorganic substance | 1358 | 43 | 10.22 | 2E-16 | 5E-13 | 4 |
| 1 | GO:0006970 | response to osmotic stress | 833 | 30 | 6.27 | 7E-13 | 1E-09 | 4 |
| 1 | GO:0042737 | drug catabolic process | 137 | 14 | 1.03 | 2E-12 | 2E-09 | 4 |
| 1 | GO:0010207 | photosystem II assembly | 157 | 14 | 1.18 | 1E-11 | 1E-08 | 4 |
| 1 | GO:0009657 | plastid organization | 388 | 21 | 2.92 | 3E-11 | 2E-08 | 4 |
| 1 | GO:0006364 | rRNA processing | 244 | 15 | 1.84 | 5E-10 | 3E-07 | 4 |
| 1 | GO:0007030 | Golgi organization | 177 | 13 | 1.33 | 9E-10 | 5E-07 | 4 |
| 1 | GO:0019682 | glyceraldehyde-3-phosphate metabolic pro... | 334 | 16 | 2.51 | 5E-09 | 2E-06 | 4 |
| 1 | GO:0010114 | response to red light | 104 | 10 | 0.78 | 6E-09 | 3E-06 | 4 |
| 1 | GO:0006598 | polyamine catabolic process | 36 | 7 | 0.27 | 8E-09 | 3E-06 | 4 |
| 1 | GO:0006754 | ATP biosynthetic process | 202 | 12 | 1.52 | 4E-08 | 2E-05 | 4 |
| 1 | GO:0006833 | water transport | 141 | 10 | 1.06 | 1E-07 | 4E-05 | 4 |
| 1 | GO:0042398 | cellular modified amino acid biosyntheti... | 55 | 7 | 0.41 | 2E-07 | 5E-05 | 4 |
| 1 | GO:0009266 | response to temperature stimulus | 950 | 24 | 7.15 | 2E-07 | 5E-05 | 4 |
| 1 | GO:0006544 | glycine metabolic process | 57 | 7 | 0.43 | 2E-07 | 6E-05 | 4 |
| 1 | GO:0052541 | plant-type cell wall cellulose metabolic... | 24 | 5 | 0.18 | 9E-07 | 2E-04 | 4 |
| 1 | GO:0015977 | carbon fixation | 12 | 4 | 0.09 | 2E-06 | 4E-04 | 4 |
| 1 | GO:0019760 | glucosinolate metabolic process | 204 | 11 | 1.53 | 3E-06 | 6E-04 | 4 |
| 1 | GO:0080147 | root hair cell development | 203 | 10 | 1.53 | 3E-06 | 7E-04 | 4 |
| 1 | GO:0015706 | nitrate transport | 206 | 10 | 1.55 | 4E-06 | 8E-04 | 4 |
| 1 | GO:0006006 | glucose metabolic process | 256 | 11 | 1.93 | 4E-06 | 8E-04 | 4 |
| 1 | GO:0042254 | ribosome biogenesis | 339 | 23 | 1.22 | 7E-19 | 3E-15 | 6 |
| 1 | GO:0009220 | pyrimidine ribonucleotide biosynthetic p... | 133 | 9 | 0.48 | 1E-09 | 3E-06 | 6 |
| 1 | GO:0046686 | response to cadmium ion | 466 | 12 | 1.68 | 1E-07 | 1E-04 | 6 |
| 1 | GO:0006096 | glycolytic process | 194 | 8 | 0.7 | 5E-07 | 5E-04 | 6 |
| 2 | GO:0009408 | response to heat | 297 | 23 | 1.81 | 4E-19 | 2E-15 | 1 |
| 2 | GO:0009644 | response to high light intensity | 223 | 19 | 1.36 | 1E-16 | 2E-13 | 1 |
| 2 | GO:0042542 | response to hydrogen peroxide | 194 | 17 | 1.18 | 3E-15 | 4E-12 | 1 |
| 2 | GO:0006457 | protein folding | 288 | 16 | 1.75 | 2E-11 | 3E-08 | 1 |
| 2 | GO:0010876 | lipid localization | 253 | 11 | 1.54 | 4E-07 | 4E-04 | 1 |
| 2 | GO:0048316 | seed development | 714 | 34 | 5.51 | 9E-18 | 4E-14 | 2 |
| 2 | GO:0019915 | lipid storage | 102 | 15 | 0.79 | 2E-15 | 4E-12 | 2 |
| 2 | GO:0050826 | response to freezing | 96 | 13 | 0.74 | 5E-13 | 7E-10 | 2 |
| 2 | GO:0090351 | seedling development | 245 | 15 | 1.89 | 6E-10 | 6E-07 | 2 |
| 2 | GO:0009415 | response to water | 423 | 18 | 3.27 | 5E-09 | 4E-06 | 2 |
| 2 | GO:0010182 | sugar mediated signaling pathway | 113 | 9 | 0.87 | 2E-07 | 2E-04 | 2 |
| 2 | GO:0097305 | response to alcohol | 611 | 19 | 4.72 | 4E-07 | 2E-04 | 2 |
| 2 | GO:0016114 | terpenoid biosynthetic process | 262 | 16 | 2.02 | 6E-07 | 3E-04 | 2 |
| 3 | GO:0009062 | fatty acid catabolic process | 218 | 9 | 0.64 | 1E-08 | 6E-05 | 5 |

**Supplemental Table S2.** Distant eQTL hotspots of the four seed germination stages. These hotspots were identified by dividing the genome into bins of 2 Mbp and performing a test to determine whether the number of distant eQTLs on a particular bin is higher than expected ($p > 0.0001$) assuming a Poisson distribution. Seed germination phenotype and metabolite data were taken from Joosen et al. (2012) and Joosen et al. (2013), respectively. Abbreviations in the phenotypes are as follows: AR, after-ripened; CD, controlled deterioration; NS, no stratification; WS, with stratification; Gmax, maximum percentage of seed germination; t10, time required for 10% of seeds to germinate; t50, time required for 50% of seeds to germinate; U8416, uniformity of germination that is the time interval between 84% and 16% of seeds to germinate; AUC, the area under the curve that is a parameter that combines Gmax, t50, and U8416. More information about the phenotypes can be seen in Joosen et al. (2012).

| hotspot ID | position | distant eQTLs | enriched GO terms[1] | metabolite | phenotype* |
|---|---|---|---|---|---|
| **PD1** | ch1:6-10Mb | 43 | negative regulation of seed germination, alpha-ketoglutarate transport, oxaloacetate transport, RNA secondary structure unwinding, medium-chain fatty acid metabolic process, gibberellin biosynthetic process, alternative mRNA splicing via spliceosome, ammonia assimilation cycle, malate transmembrane transport, response to carbon dioxide, regulation of seed dormancy process | Sinapate | Stratification.AR.AUC, Stratification.AR.t50, ABA.NS.Gmax, Cold.NS.Fresh.t10 |
| **PD2** | ch3:8-12Mb | 69 | positive regulation of protein export from nucleus, detection of brassinosteroid stimulus, galactose catabolic process | two unknown metabolites | Heat.NS.AR.t50 |
| **AR1** | ch2:12-14Mb | 16 | none | none | none |
| **AR2** | ch3:2-4Mb | 20 | response to oxidative stress, response to light intensity, translation, phospholipid transfer to membrane, proline catabolic process, asparagine biosynthetic process, glutamate biosynthetic process, protein folding, response to heat | Sinapate | Heat.NS.AR.Gmax, Heat.NS.AR.AUC, Heat.WS.AR.AUC |
| **IM1** | ch5:6-8Mb | 19 | Lewis a epitope biosynthetic process, DNA catabolic process | D-maltose and 23 unknown metabolites | NaCl.NS.t10 |
| **IM2** | ch5:22-26Mb | 69 | negative regulation of chromatin silencing, seed germination, response to karrikin, entrainment of circadian clock by photoperiod, proline catabolic process, glutamate biosynthetic process | L-isoleucine, pyruvic acid, 2-hydroxybutyric acid, alanine, urea, oxoglutaric acid | NaCl.NS.Gmax, NaCl.WS.U8416, Mannitol.WS.AUC, CD.NS.AR.Gmax, Stratification.AR.Gmax, Mannitol.WS.t50, Stratification.AR.U8416, Stratification.Fresh.Gmax, Stratification.Fresh.AUC, Mannitol.NS.t10, Mannitol.NS.U8416, ABA.NS.AUC, Stratification.Fresh.t50, ABA.NS.Gmax, ABA.NS.t50, AfterRipening.t10, AfterRipening.t50, Stratification.Fresh.t10, |

| | | | | | |
|---|---|---|---|---|---|
| | | | | | Heat.NS.AR.AUC, ABA.NS.t10, ABA.WS.Gmax, ABA.WS.AUC, ABA.WS.t10, ABA.WS.t50, ABA.WS.U8416, Size.imbibed.seed.Area, Mannitol.NS.t50, Heat.WS.AR.U8416, AR.Gmax.ns, Heat.NS.AR.Gmax, Cold.NS.Fresh.Gmax |
| **RP1** | ch1:0-2Mb | 23 | medium-chain fatty acid metabolic process | none | AR.U8416.ns |
| **RP2** | ch1:6-8Mb | 18 | none | none | Stratification.AR.AUC, Stratification.AR.t50, ABA.NS.Gmax |
| **RP3** | ch5:14-16Mb | 21 | response to karrikin, pigment biosynthetic process, response to far red light, response to UV-B, response to red light, positive regulation of catalytic activity, response to blue light, flavonoid biosynthetic process, maltose metabolic process, pentose-phosphate shunt, starch biosynthetic process, response to sucrose, photosynthetic electron transport in photosystem II, membrane disassembly, rRNA processing, cellular response to calcium ion, photosynthesis, light harvesting in photosystem II, fructose 1,6-bisphosphate metabolic process, fructose metabolic process, cellular response to UV-A, photoprotection, cellular response to high light intensity, phenylpropanoid biosynthetic process, monosaccharide transmembrane transport, cytidine deamination, regulation of stomatal closure, photosystem II assembly, de-etiolation | none | CD.NS.AR.t50 |
| **RP4** | ch5:24-26Mb | 96 | seed development, lipid storage, response to acid chemical, seed germination, response to inorganic substance, response to freezing, toxin metabolic process, response to cyclopentenone, response to osmotic stress, defense response to fungus incompatible interaction, response to hypoxia, sugar mediated signaling pathway, abscission, protein homooligomerization, fatty acid catabolic process, leaf senescence, D-serine biosynthetic process, inorganic anion transport | Alanine, fumaric acid, glycine, L-aspartic acid, L-glutamine, L-lysine, L-methionine, L-phenylalanine, L-serine, L-tyrosine, L-valine, malic acid, sucrose, and other 7 unknown metabolites | Stratification.AR.U8416, Stratification.Fresh.Gmax, Stratification.Fresh.AUC, Mannitol.NS.t10, Mannitol.NS.U8416, ABA.NS.AUC, Stratification.Fresh.t50, ABA.NS.Gmax, ABA.NS.t50, AfterRipening.t10, AfterRipening.t50, Stratification.Fresh.t10, Heat.NS.AR.AUC, ABA.NS.t10, ABA.WS.Gmax, ABA.WS.AUC, ABA.WS.t10, ABA.WS.t50, |

ABA.WS.U8416, Size.imbibed.seed.Area, Mannitol.NS.t50, Heat.WS.AR.U8416, AR.Gmax.ns, Heat.NS.AR.Gmax, Cold.NS.Fresh.Gmax

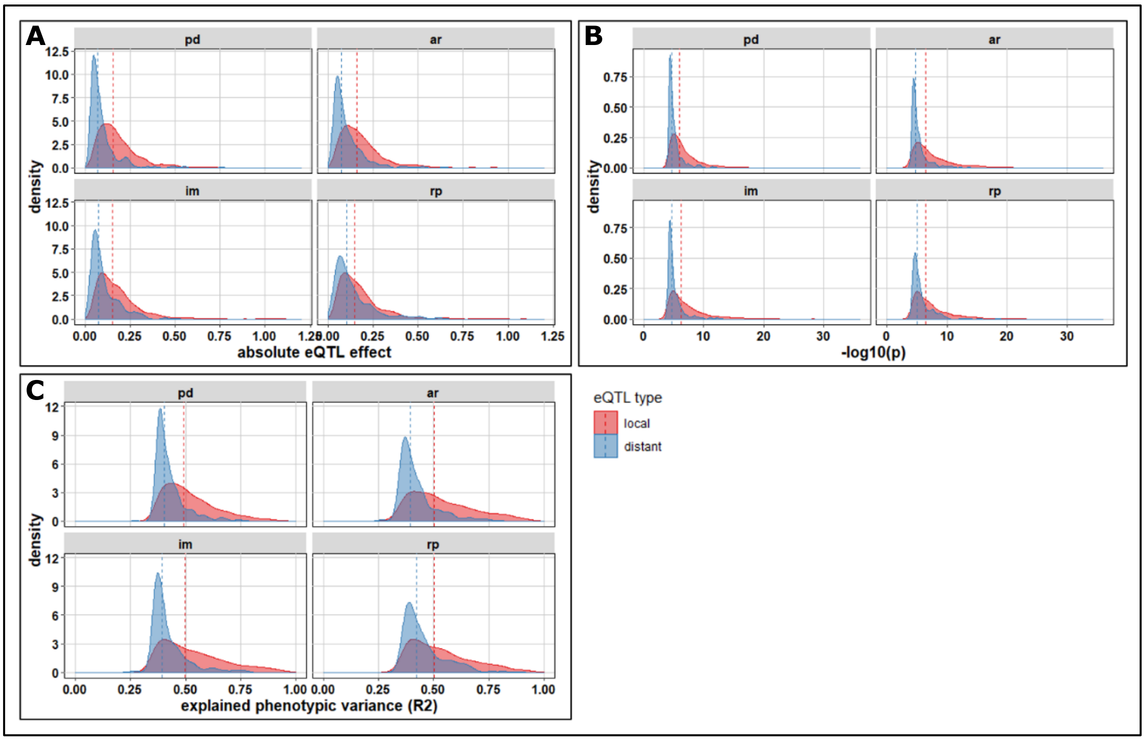

**Figure S1.** Density distribution of the absolute eQTL effect, -log(p), and explained phenotypic variance ($R^2$) for local and distant eQTLs. The eQTL effect is the estimated effect of an eQTL to the log intensity of the gene transcript, and the significance of this estimation is reflected in the -log10(p) value. Meanwhile, the value of $R^2$ represents the fraction of the total variation that is explained by the simple linear regression model.

**Supplemental Table S3**. The mean rank and standard deviation of candidate genes as the most likely causal genes for the RP4 hotspot across different thresholds.

| candidate genes | mean rank | SD rank |
|---|---|---|
| AT5G61590 (DEWAX) | 1.89 | 0.71 |
| AT5G62490 | 2.15 | 1.65 |
| AT5G62540 | 2.24 | 0.9 |
| AT5G59690 | 4.16 | 0.52 |
| AT5G63020 | 6.43 | 1.74 |
| AT5G63030 | 6.6 | 2.07 |
| AT5G63150 | 7.47 | 2.79 |
| AT5G64510 | 9.82 | 3.78 |
| AT5G64310 | 11.18 | 3.05 |
| AT5G65060 | 11.36 | 3.69 |
| AT5G65020 | 11.53 | 3.16 |
| AT5G60370 | 11.79 | 2.3 |
| AT5G62340 | 12.37 | 1.94 |
| AT5G61010 | 13.08 | 3.45 |
| AT5G61190 | 14.83 | 2.5 |

| | | |
|---|---|---|
| AT5G64120 | 15.05 | 1.74 |
| AT5G59680 | 15.47 | 3.91 |
| AT5G61290 | 16.83 | 4.04 |
| AT5G63087 | 17.84 | 1.62 |
| AT5G62420 | 17.91 | 2.36 |

**Supplemental Table S4**. The mean rank and standard deviation of candidate genes as the most likely causal genes for the PD2 hotspot across different thresholds.

| candidates | mean rank | SD rank |
|---|---|---|
| AT3G26744 (ICE1) | 1.38 | 2.74 |
| AT3G27350 | 3.63 | 4.24 |
| AT3G27610 | 4.05 | 1.15 |
| AT3G27925 | 4.36 | 5.59 |
| AT3G26950 | 5.09 | 2.35 |
| AT3G27280 | 5.87 | 2.48 |
| AT3G27360 | 7.21 | 3.11 |
| AT3G24550 | 11.15 | 7.55 |
| AT3G27930 | 11.72 | 3.31 |
| AT3G24503 | 12.81 | 7.19 |
| AT3G25160 | 13.95 | 5.73 |
| AT3G27220 | 14.06 | 6.79 |
| AT3G27430 | 14.52 | 6.12 |
| AT3G24315 | 14.84 | 2.76 |
| AT3G28345 | 14.85 | 4.31 |
| AT3G26240 | 15.41 | 3.13 |
| AT3G29090 | 15.79 | 5.51 |
| AT3G28060 | 19.61 | 4.4 |
| AT3G28460 | 20.35 | 7.21 |
| AT3G25870 | 20.65 | 2.73 |
| AT3G24830 | 20.9 | 2.77 |
| AT3G28150 | 21.11 | 5.32 |
| AT3G24030 | 22.44 | 3.66 |
| AT3G27670 | 24.43 | 4.04 |
| AT3G22890 | 25.92 | 4.05 |
| AT3G23090 | 26.67 | 4.11 |
| AT3G25040 | 27.8 | 5.23 |
| AT3G26170 | 29.02 | 5.79 |
| AT3G27440 | 29.5 | 8.28 |
| AT3G28700 | 30.01 | 5.18 |
| AT3G24255 | 31.31 | 5.21 |
| AT3G28956 | 31.32 | 5.39 |

| | | |
|---|---|---|
| AT3G28940 | 31.67 | 5.16 |
| AT3G27660 | 32.19 | 4.47 |
| AT3G24070 | 33.18 | 6.39 |
| AT3G27560 | 33.86 | 4.48 |
| AT3G23030 | 33.97 | 5.75 |
| AT3G26380 | 36.79 | 7.84 |
| AT3G24630 | 37.36 | 4.34 |
| AT3G28740 | 37.96 | 7.65 |
| AT3G24430 | 38.98 | 5.73 |
| AT3G25780 | 39.89 | 3.91 |
| AT3G25013 | 40.35 | 4.7 |
| AT3G28920 | 42.07 | 1.73 |
| AT3G29075 | 45 | 0.03 |

**Supplemental Table S5**. The location and type of SNPs on candidate genes for the RP4 eQTL hotspot and *MUM2*

| TAIR ID | SNP location/type | | | |
|---|---|---|---|---|
| | 5' UTR | non-synonymous | synonymous | 3' UTR |
| AT5G61590 (DEWAX) | 0 | 2 | 4 | 2 |
| AT5G63800 (MUM2) | 0 | 2 | 2 | 2 |
| AT5G62490 | 0 | 0 | 0 | 1 |
| AT5G62540 | 4 | 1 | 1 | 1 |
| AT5G59690 | 1 | 1 | 1 | 0 |
| AT5G63020 | 0 | 14 | 11 | 2 |
| AT5G63030 | 1 | 0 | 1 | 1 |
| AT5G63150 | 0 | 1 | 1 | 2 |
| AT5G64510 | 0 | 1 | 1 | 0 |
| AT5G64310 | 1 | 0 | 1 | 2 |
| AT5G65060 | 0 | 0 | 0 | 1 |
| AT5G65020 | 1 | 0 | 0 | 0 |
| AT5G60370 | 1 | 3 | 3 | 2 |
| AT5G62340 | 0 | 2 | 2 | 1 |
| AT5G61010 | 3 | 0 | 0 | 1 |
| AT5G61190 | 0 | 10 | 7 | 2 |
| AT5G64120 | 0 | 0 | 1 | 0 |
| AT5G59680 | 0 | 7 | 18 | 0 |
| AT5G61290 | 0 | 1 | 0 | 0 |
| AT5G63087 | 1 | 2 | 0 | 2 |
| AT5G62420 | 0 | 1 | 12 | 0 |

**Supplemental Table S6**. The location and type of SNPs on candidate genes for the PD2 eQTL hotspot.

| TAIR ID | SNP location/type | | | |
|---|---|---|---|---|
| | 5' UTR | non-synonymous | synonymous | 3' UTR |
| AT3G26744 (ICE1) | 0 | 1 | 14 | 1 |
| AT3G27350 | 6 | 5 | 10 | 6 |
| AT3G27610 | 0 | 5 | 7 | 1 |
| AT3G27925 | 0 | 3 | 11 | 0 |
| AT3G26950 | 0 | 2 | 0 | 0 |
| AT3G27280 | 0 | 0 | 0 | 2 |
| AT3G27360 | 0 | 0 | 3 | 6 |
| AT3G24550 | 0 | 4 | 7 | 1 |
| AT3G27930 | 1 | 1 | 3 | 3 |
| AT3G24503 | 0 | 0 | 12 | 0 |
| AT3G25160 | 0 | 1 | 3 | 1 |
| AT3G27220 | 2 | 3 | 2 | 0 |
| AT3G27430 | 4 | 1 | 7 | 4 |
| AT3G24315 | 0 | 0 | 0 | 0 |
| AT3G28345 | 0 | 3 | 6 | 1 |
| AT3G26240 | 4 | 30 | 37 | 2 |
| AT3G29090 | 1 | 0 | 10 | 1 |
| AT3G28060 | 0 | 6 | 0 | 0 |
| AT3G28460 | 0 | 0 | 0 | 0 |
| AT3G25870 | 0 | 1 | 1 | 0 |
| AT3G24830 | 0 | 0 | 1 | 2 |
| AT3G28150 | 0 | 8 | 12 | 3 |
| AT3G24030 | 0 | 0 | 0 | 4 |
| AT3G27670 | 0 | 17 | 6 | 0 |
| AT3G22890 | 0 | 0 | 1 | 1 |
| AT3G23090 | 1 | 0 | 0 | 1 |
| AT3G25040 | 1 | 1 | 5 | 2 |
| AT3G26170 | 0 | 0 | 1 | 0 |
| AT3G27440 | 0 | 0 | 8 | 0 |
| AT3G28700 | 0 | 1 | 0 | 0 |
| AT3G24255 | 0 | 0 | 0 | 0 |
| AT3G28956 | 0 | 1 | 0 | 0 |
| AT3G28940 | 0 | 4 | 2 | 7 |
| AT3G27660 | 1 | 0 | 3 | 0 |
| AT3G24070 | 2 | 1 | 0 | 0 |
| AT3G27560 | 3 | 1 | 11 | 4 |
| AT3G23030 | 0 | 1 | 3 | 1 |
| AT3G26380 | 0 | 5 | 10 | 1 |
| AT3G24630 | 0 | 2 | 1 | 0 |
| AT3G28740 | 0 | 1 | 3 | 0 |
| AT3G24430 | 0 | 0 | 0 | 0 |

| | | | | |
|---|---|---|---|---|
| AT3G25780 | 0 | 1 | 0 | 1 |
| AT3G25013 | 0 | 2 | 0 | 0 |
| AT3G28920 | 1 | 1 | 2 | 0 |
| AT3G29075 | 1 | 2 | 4 | 1 |

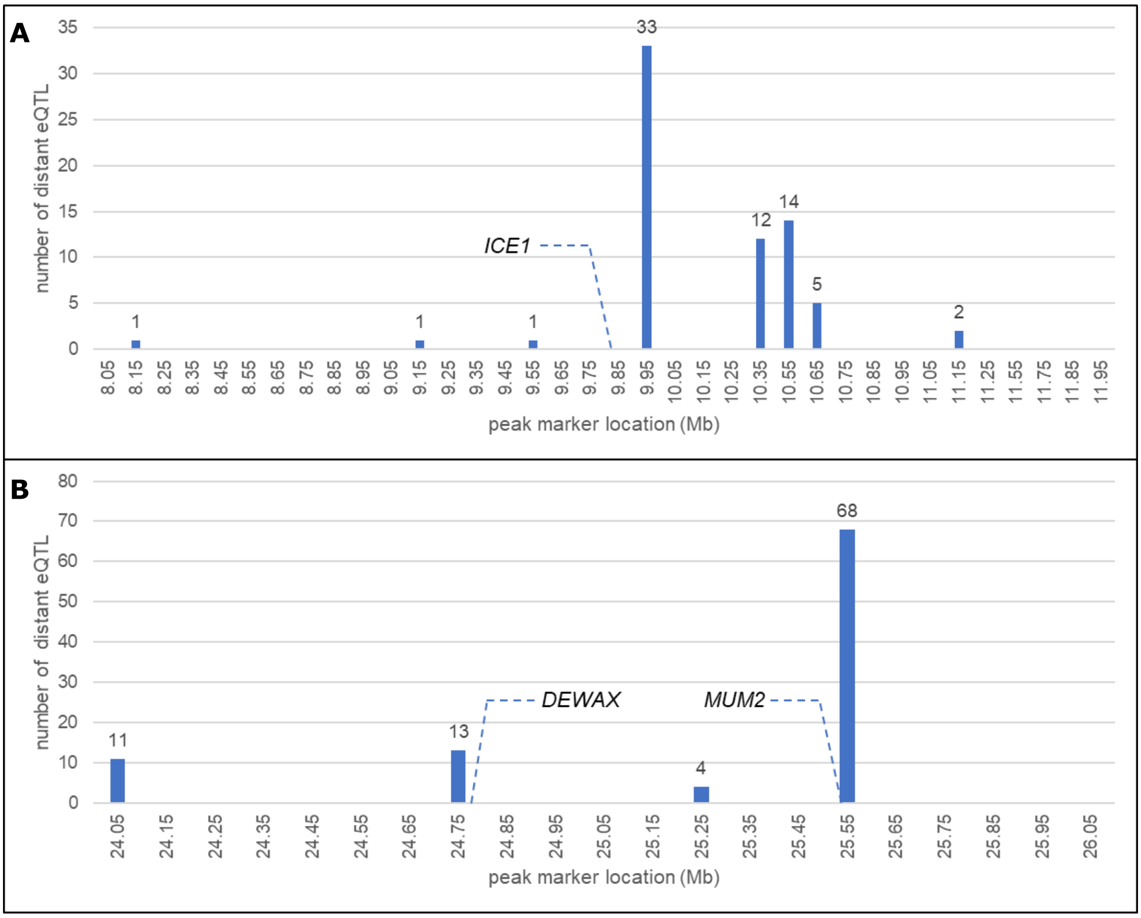

**Figure S2.** The histogram of the number of distant eQTLs per marker location for the PD2 (A) and RP4 (B) hotspot.
